# Signal to Noise Ratio as a Cross-Platform Metric for Intraoperative Fluorescence Imaging

**DOI:** 10.1101/847863

**Authors:** Asmaysinh Gharia, Efthymios P. Papageorgiou, Simeon Giverts, Catherine Park, Mekhail Anwar

## Abstract

Real-time molecular imaging to guide curative cancer surgeries is critical to ensure removal of all tumor cells, however visualization of microscopic tumor foci remains challenging. Wide variation in both imager instrumentation and molecular labeling agents demands a common metric conveying the ability of a system to identify tumor cells. Microscopic disease, comprised of a small number of tumor cells, has a signal on par with the background, making the use of signal (or tumor) to background ratio inapplicable in this critical regime. Therefore, a metric that incorporates the ability to subtract out background, evaluating the signal itself relative to the sources of uncertainty, or noise is required. Here we introduce the signal-to-noise ratio (SNR) to characterize the ultimate sensitivity of an imaging system, and optimize factors such as pixel size. Variation in the background (noise) are due to electronic sources, optical sources, and spatial sources (heterogeneity in tumor marker expression, fluorophore binding, diffusion). Here we investigate the impact of these noise sources and ways to limit its effect on SNR. We use empirical tumor and noise measurements to procedurally generate tumor images and run a monte carlo simulation of microscopic disease imaging to optimize parameters such as pixel size.

## Introduction

Knowledge of the presence of tumor cells is essential for cancer surgery. Small numbers of tumor cells, impossible to detect with the unaided eye or by touch, are often left behind, leading to positive margins that are strikingly common. Positive margins occur in 25-40% of breast cancer surgeries ^1,2^ and 20-50% of high risk prostate cancer surgeries (Zeitman et al. 1993). Positive margins, or microscopic residual disease (MRD) are consequential, significantly increasing the risk that cancer returns across cancer types, for example doubling the recurrence in breast cancer leading to decreased survival ^3^. Similarly, MRD in prostate cancer increases the risk of recurrence 2-4 times ^4–7^. Efforts to address MRD have long centered around physician judgement through preoperative imaging, and intraoperative sight and touch. However these techniques are limited to millimeter to centimeter scale resolution - equivalent to 10^4^-10^9^ cells, orders of magnitude above the needed threshold of detection to ensure a margin negative outcome. Gold-standard methods of margin detection rely on pathologic examination of the excised specimen, and if the specimen surface includes tumor cells (called a positive margin), additional therapy is performed at a later date - for example re-excision for breast cancer, and post-operative radiation for prostate cancer.

Current strategies for intraoperative tumor identification face challenges when assessing microscopic disease. Intraoperative specimen radiography can assist with verification of gross tumor removal, but not for portions of the tumor remaining in the patient. Strategies for identifying tumor margin have focused on frozen section and touch prep analysis. Frozen section analysis, particularly challenging with the fatty tissue in breast cancer, is hindered by false negatives (Riedl, Fitzal et al. 2009), requires a pathologist present at the time of surgery, limited in the area that can be evaluated potentially missing disease, and significantly prolongs operative time.

Real-time molecular imaging on the other hand, offers the opportunity to visualize MRD intraoperatively, directly in the tumor bed, enabling treatment of all disease at the time of the initial operation. Consequently, the need for imaging microscopic disease has driven the development of highly sensitive intraoperative imagers. Taking advantage of the growing armamentarium of fluorescently-tagged molecular imaging agents, fluorescence imaging has moved to the forefront of intraoperative visualization techniques (Frangioni 2008). While a wide range of intraoperative imagers exist, no standardized metric exists to evaluate their performance, particularly when coupled to a targeted molecular imaging agent. Therefore, a platform *independent* method is needed to quantify the ability of imagers to detect microscopic disease intraoperatively.

The default method for identifying residual tumor using intraoperative imagers has been physician identification from an image. Efforts to define a common quantification metric for imaging tools have centered around the signal-to-background ratio (SBR) ^8,9^. Implicit in this metric is a tumor signal significantly above background - true for larger tumor foci, but not necessarily for microscopic disease, which is often just above background contributed by non-specific binding, autofluorescence, and other optical and electronic sources. However, to properly identify microscopic tumor foci in an image, the background must be accurately subtracted - often in software - using a combination of background subtraction and image recognition to achieve sensitivities far beyond human visual identification. This makes accurate determination of background critical, as any error in background estimation translates directly into an error in signal.

The background variation in biological systems can be analogized to measurement uncertainty in general, often called “noise”, and for images is quantified as spatial noise. When combined with signal intensity, this leads to a quantifiable signal to noise ratio for detecting microscopic disease in an imaging system. Here we propose signal-to-noise ratio (SNR) as a figure of merit for optical detection of microscopic disease, which represents the fundamental limit of electronic and computer aided detection. This metric can be used to compare sensitivity across imaging systems and define the ultimate limits of detection for a system.

To quantify SNR, we measure both the signal and background as well as their variation. The signal is defined as the number of photons collected from a tumor foci, and this paper addresses the identification and quantification of noise sources in the imaging system such as electronic noise and spatial noise. Key to accurate background subtraction (as shown in Figure 1), these factors are affected by the detector sensitivity, optical background rejection, properties of the imaging marker (antibody binding kinetics), antigen expression by the tumor and normal cells, and pixel size. The latter parameter is critical, as smaller pixels (higher resolution) are not always “better” - too small of a pixel may only sample noise with minimal signal, while too large of a pixel may washout tumor signal by averaging with background. Conversely, a pixel size larger than a single cell is still capable of single cell detection if the background is accurately subtracted. Thus, maximum SNR is intrinsically linked to pixel size.

**Figure 1.**
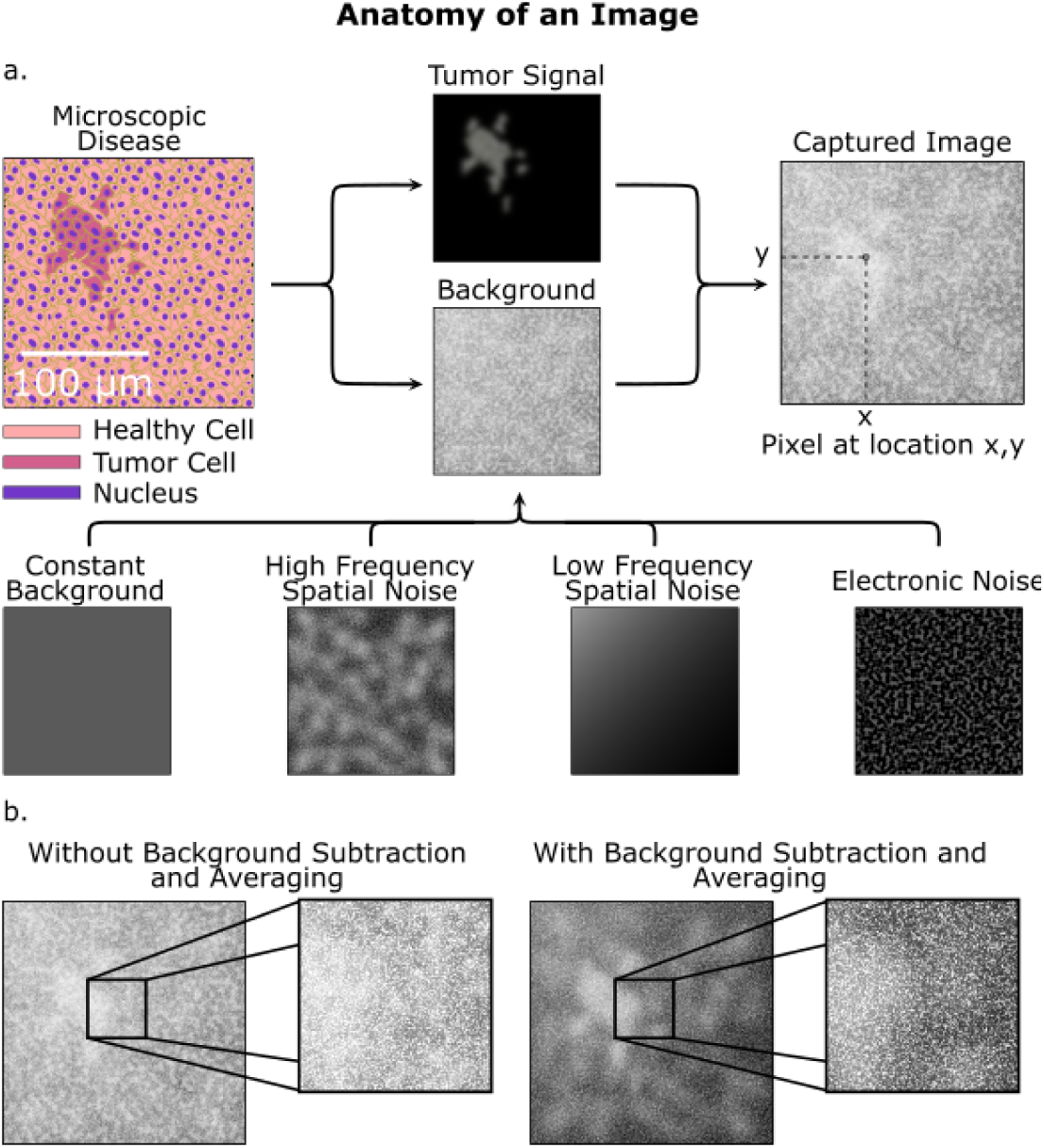
Sources of signal, background and noise. **(a)** A simulated image of microscopic disease including background and noise sources that obfuscate the tumor signal. Both the tumor area and background are procedurally generated. **(b)** Without background subtraction and averaging, the tumor is hard to identify, while with background subtraction the tumor is more apparent.

Of the noise sources, this paper focuses on defining and quantifying spatial noise so that an accurate SNR is defined. Electronic (e.g. time varying) noise can be mitigated with sufficient image averaging (or equivalently, longer integration times), but spatial noise arising from variations in the underlying tissue and staining conditions cannot, thus driving a need to study the impact of spatial noise on the SNR for background subtraction. Analogous to time-varying noise sources, spatial noise is composed of both high and low frequency components. Similar in concept to averaging or longer integration time for time-varying noise, high-frequency spatial noise can be reduced by imaging a larger area for each pixel (e.g. larger pixel size); however, this comes at the expense of spatial resolution, and can lead to errors by integrating large fluctuations in slowly varying background intensity. Therefore, there is an ideal pixel size to optimize SNR for a given imaging system.

In this paper, we outline a general method for evaluating the SNR of any optical imager in combination with a biologic labelling of tumor cells, and relate this to optimal pixel size. Since the analysis is based on the image itself, this method can be used as a platform independent metric to compare imagers (and biologics) in the evaluation of microscopic disease, essential for modern optical surgical navigation. Quantification of imaging systems using SNR allows single cell imaging, even with systems whose spatial resolution is below that of a single cell. We discuss the various contributions to the tumor signal and the sources of background and their inherent variability, which contributes noise as shown in Table 1.

**Table 1.**

| ***Signal Sources*** | ***Background Sources*** | ***Noise Sources*** |
| --- | --- | --- |
| *Number tumor cells* | *Dark Current* | *Electronic (Shot Noise)* |
| *Molecular labelling specificity* | *Non-specific binding* | *Cell to cell variability* |
| *Molecular label concentration* | *Healthy cell antigen expression* | *Diffusion* |
| *Illumination intensity* | *Optical bleed through* | *Tissue heterogeneity* |
| *Fluorophore Quantum Efficiency* | *Autofluorescence* |  |
| *Electronic Responsivity* |  |  |
| *Pixel Size* |  |  |

To illustrate our methodology, we quantified spatial noise with immunofluorescence imaging of breast and prostate cancer cells, using both *in vitro* and *in vivo* molecular staining. For the purposes of molecular labeling, we used a model system of HER2-overexpressing (HER2+) breast cancer cell lines (SKBR3, HCC1569) compared against HER2-negative (HER2-) cell lines (S1, MDA-MB-231) with trastuzumab ^10^, an antibody targeting the HER2 receptor. Similarly we use PMSA-overexpressing prostate cancer cell lines (LnCAP) and PSMA-negative (PC3) with J591, a humanized antibody against PSMA ^11–14^. As a demonstration of this technique, we quantified the signal from tumor, and the sources of noise in Table 1, with a fluorescence microscope. Figure 1 illustrates these sources of noise, drawing a distinction between high frequency spatial noise and low frequency spatial noise. High frequency spatial noise, which varies rapidly over the image - consists of variations in antibody binding per cell and natural tissue (and tumor) heterogeneity.

Low frequency spatial noise - which varies slowly over the entire image - can be a result of gradients in antibody concentration due to diffusion and tumor perfusion and vascularity. Using these measured metrics, we then randomly generate images of clusters of cells of specific size, and quantify the SNR across varying arrangements of cell clusters. We use the characterization data obtained from real cell samples to make a simulated model of residual cancer tissue in order to determine the optimal pixel size for an intraoperative imager. The method and algorithm can be easily generalized for any cell line and antibody combination.

## Methods

### Cell Culture

#### Breast Cancer Cell Lines

*In Vitro* breast cancer cell cultures consisted of SKBR3 (HER2-overexpressing) and S1 (HER2-negative) (from ATCC) cultured in RPMI with 10% FBS. *Prostate Cancer Cell Lines*: Prostate cell cultures consisted of LnCAP cells (PSMA overexpressing) and PC3 cells (PSMA negative).

### In Vivo Mouse Models

To determine the *in vivo* kinetic and spatial distribution of trastuzumab, we subcutaneously implanted HER2 over-expressing HCC1569 cells and MDA-MB-231 (HER2-negative) cells as a negative control in nude mice, and injected increasing amounts of trastuzumab via intraperitoneal injection. Mice were sacrificed at 24, 48 and 96 hours and tumor, kidney, muscle, and liver were removed and stained for trastuzumab binding.

### Staining

#### Fixation

Mouse tissue sections were fixed with 2% Paraformaldehyde in phosphate-buffered saline (PBS) solution for 20 minutes at room temperature. Slides are then washed with PBS and glycine solution.

#### Blocking

Tissue sections were blocked with 10% Goat Serum in IF buffer.

#### Immunostaining

Sections were then further stained with the nuclear stain DAPI to simplify locating cells using the microscope. Mounting: Coverslips were then mounted with Vectashield Storage Medium (Vector Labs H1000)

### Imaging Procedure

Images were taken with Leica DMIRB with 20X objective and standard FITC filter sets (Chroma) using a Hamamatsu ORCA-Flash4.0 V2 camera. Tissue images, used for *in vivo* binding quantification were taken from the center of tissue samples. Background variation data were taken by imaging 66 μm x 66 μm areas, then shifting the slide by 59.4 μm and taking another image. This provided a 10% overlap between images, allowing for image stitching.The procedure was repeated across the entire tissue slice. The slide movement was precisely controlled using a Thorlabs XY Mechanized Stage and the stage and camera was controlled by Micro-Manager ^15^. Individual cells were identified using CellProfiler ^16^, and the total fluorescence intensity was quantified. The number of antibodies corresponding to the fluorescence intensity value was determined by imaging a set of reference dilutions of FITC conjugated secondary antibody and comparing fluorescence intensity. A linear fit was established, defining the relationship between number of antibodies and fluorescent intensity per pixel using the same objective and integration time. Using this calibration curve, the fluorescence intensity of each cell was converted to the number of antibodies bound. Diffusion across tissue was estimated using MATLAB to determine average differences in intensity across an entire tissue slice.

### Monte Carlo Simulation of SNR

We generated images of cell clusters to estimate the maximal SNR and optimal pixel size for our imaging sensor. Each image consists of a randomly generated cell clusters of ~100 cells. The cell images are procedurally generated using Perlin noise (Perlin 2002) to create a binary mask to demarcate tumor versus non-tumor areas as seen in Figure 1a (Tumor Signal). To accurately replicate both the signal and background intensity we assign cell intensity and background on the values found in Figure 3 for HER2-overexpressing and HER2-negative cells, respectively.

Background is created as a random matrix with the same average intensity and variation (quantified as the standard deviation) as non-specifically labeled cells imaged within the MDA-MB-231 (HER2-negative) tumor stained with trastuzumab. A 5%/mm intensity gradient is added to mimic the gradient measured in Figure 4. Similarly, to accurately replicate the intensity and spatial noise of the tumor signal, a random matrix is created with the same average intensity and standard deviation as specifically labeled cells in the HCC1569 (HER2-overexpressing) tumor stained with trastuzumab. This matrix is then clipped (multiplied) by the binary mask defined earlier. The background matrix and tumor matrix are then summed resulting in a simulated image where the background has the same variance and average intensity as HER2-negative tissue and the tumor areas have the same variance and intensity as HER2-overexpressing tissue.

## Results

### SNR as a Metric for Intraoperative Detection of Microscopic Disease

The SNR defines the detection limit of the complete imaging system and is defined as:

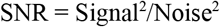

The MRD signal is defined as the total signal, or number of photons, T, received by a pixel (gathered and converted to electrons by a pixel’s photosensitive element). This consists of both the photons emitted by the optically labelled tumor cells (called the signal, S) and the background, B, at that location (x,y) as seen in Figure 1. We can write this as:

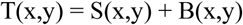

To estimate background intensity for background subtraction, we measure the pixel intensity at a location away from the microscopic tumor (x + dx, y + dy), absent of tumor (S(x+dx, y+dy) = 0), such that:

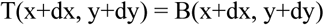

The MRD signal alone can then be estimated from these two measurements as:

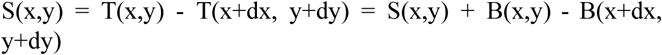

Recognizing that B(x,y) and B(x+dx, y+dy) may differ, this introduces the spatial noise in the system, and the minimum tumor signal detectable is then equal to this uncertainty, ΔB = B(x,y) - B(x+dx, y+dy), which is the noise of the system. Here we have assumed that there is sufficient averaging to reduce electronic noise to below the level of the spatial noise, and do not quantify its contribution. In this regime we find:

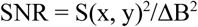

For small clusters of cells, the tumor signal is weak and is on the same order as the background intensity. We define the minimum number of detectable cells as that which gives an SNR > 10, a value ensuring the possibility of identification of tumor cells.

### Quantification of Tumor Signal

The goal of intraoperative MRD imaging is to identify, and quantify, the number of tumor cells amidst a background of physiologically similar normal tissue cells. The signal from the tumor cells is proportional to

1. The number cells to be detected (N_cell_).
2. The number of molecules labeled or bound to each cell (**α**_bound_).
3. The illumination (excitation) photon flux.
4. The fluorophore efficiency of converting those illumination photons to Stokes shifted emitted photons.

The number of antibodies labeled per cell (**α**_bound_) is a function of the tumor biomarker binding affinity, biomarker expression level and the labeling molecules exposure time to cells. The ratio of bound biomarker to tumor cells relative to healthy tissue cells is called the tumor to background ratio (TBR), used interchangeably here with SBR, and together the two quantities (TBR and **α**_bound_) can be used to quantitatively describe the biologics role in determining signal, and therefore, SNR. To demonstrate how to apply this technique, we have performed the following experiments determining TBR *in vitro* in both example breast (SKBR3, S1, HCC1569, MDA231) and prostate (LNCaP and PC3) cell lines and *in vivo* in the breast tumor model.

### In Vitro Determination of TBR and α_bound_

Labeling tumor cells *in vivo* using a molecularly targeted imaging agent is the first step ^17^ in translating the cell identifying procedures from the pathology laboratory into the real-time operating room environment. Identification of small foci (<200 cells) of fluorescently labeled MRD requires (1) accurate detection of the tumor focus and (2) differentiation of the tumor from the surrounding background, which can overwhelm and mask the small MRD signal. Here we quantify the binding of trastuzumab to HER2 overexpressing cell lines and J591 to PSMA overexpressing cell lines as model systems.

We quantify the number of fluorophore-labeled antibodies bound per tumor cell (**α_bound_**) and the relative background signal with the tumor-to-background ratio, TBR. Figure 2 illustrates *in vitro* quantification of HER2 labeling with increasing concentrations of trastuzumab. At 10 μg/ml of trastuzumab, SKBR3 cells bind ~30,000 antibodies/μm^2^ (α_bound_=3.6×10^6^ / cell), while only 1,700 antibodies/μm^2^ (8.5×10^3^ / cell) bind to S1 cells, for a TBR of 17. Higher concentrations of Trastuzumab saturate binding at 5×10^4^ antibodies/cell, although TBR is reduced due to increased background. Similar analysis on PSMA overexpressing prostate cancer cells, demonstrates an α_bound_=3.7×10^4^ / cell with a TBR of 28, consistent with the lower expression level of PSMA ^18^ versus HER2 ^19^.

**Figure 2.**
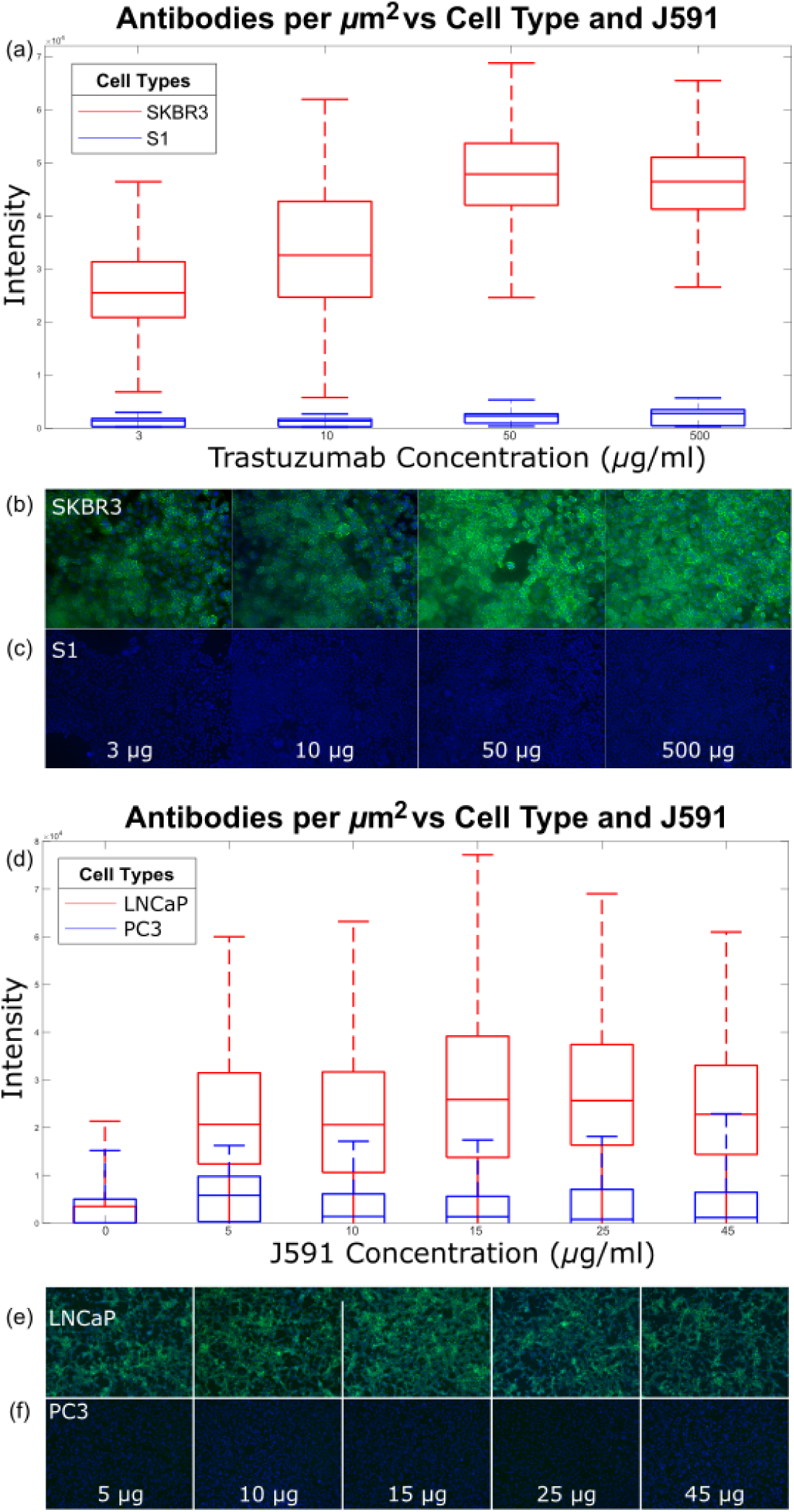
Quantification of signal and noise *in vitro*. **(a)** Quantification of trastuzumab binding to SKBR3 (HER2+) cells shows average of 30-50,000 antibodies/μm^2^, while S1 (HER2-) cells are 17X less. **(b)** Cell staining of SKBR3 and **(c)** Cell staining of S1 with trastuzumab (green = trastuzumab, blue = nucleus) with various concentrations of anti-HER2 antibody. (d-f) Similar experiment with prostate cancer cell line LNCaP (PSMA+) and PC3 (PSMA-). **(d)** Quantification of J591 binding to LNCaP cells shows 40,000 antibodies/cell. **(e,f)** Cell staining with J591 (green = J591, blue = nucleus) with various concentrations of J591 with LNCaP and PC3 respectively.

### In Vivo determination of TBR and α_bound_

To drive maximal signal for *in vivo* imaging, it is important to determine the optimal timing and concentration of a systemically injected imaging agent to maximize tumor binding (α_bound_). Studies ^20–24^ demonstrate maximal TBR 24-72 hours after injection. To determine the *in vivo* kinetic and spatial distribution of trastuzumab, we subcutaneously implanted HER2 over-expressing HCC1569 cells and MDA-MB-231 (HER2-negative) cells as a negative control in nude mice, and injected increasing amounts of trastuzumab via intraperitoneal injection. Figure 3 shows selective binding of 1mg trastuzumab to HCC1569 cells *in vivo* (~40,000 antibodies/ μm^2^, α_bound_=5×10^6^ / cell), with optimal TBR (30) at 48 hours post-injection. TBR *in vivo* exceeds *in vitro* due to receptor-mediated endocytosis of trastuzumab ^25^.

**Figure 3.**
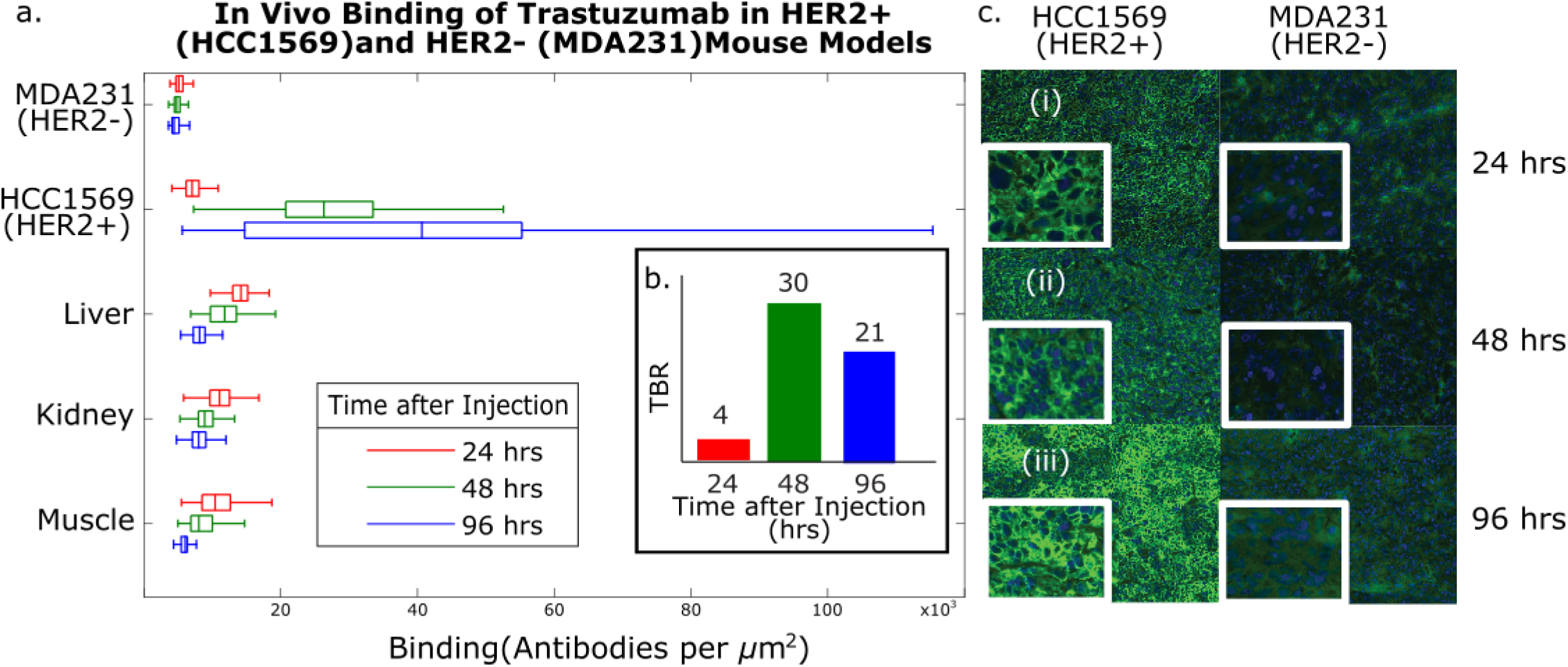
Quantification of signal and noise *in vivo*. **(a)** binding of 1mg of trastuzumab to HER2+ (HCC1569) and HER2-(triple negative, MDA-MB-231) tumors in nude mice versus time (24,48,96 hrs), stained with anti-human FITC (green) and Dapi (blue) nuclear counterstain. Binding to HER2+ cells increases with time. **(b)** TBR is 4, 30, and 21 at 24, 48 and 96 hrs post-injection. **(c)** Representative images of tumor tissue are shown at 24, 48, and 96 hours in inset (i-iii).

These experiments show how to quantify TBR, capturing both the ratio of the biological labels (e.g. antibodies) per tumor and normal tissue cell and the amount of labelling per cell. Signal is the difference between the intensity of the tumor cells and background cells. Spatial noise on the other hand is partially composed of the variation in the amount of labelling per cell. Their ratio, computed from the image, is an integrated function of both antibody specificity and imager performance, and is the key driver of detection sensitivity, governing both the signal intensity, and the background intensity. This quantitative description of biologic performance is agnostic to the imaging instrument itself (relying only on the final image), and as such can both be used to predict how sensitivity will vary with changing TBR and to compare sensitivity across imaging systems.

Additional determinants of the tumor signal include factors such as illumination intensity, fluorophore quantum efficiency, photon gathering, and elements of electronic detection such as pixel responsivity, pixel capacity, and integration time. Here we qualitatively describe their impact on the signal.

### Illumination Intensity

Illumination intensity proportionally increases the intensity of the fluorescent signal and the optical sources of background. However, given a fixed total imaging time to evaluate SNR, increasing illumination decreases electronic noise as multiple images can be averaged in the same period due to the increased number of photons reaching the sensor per unit time (decreasing the integration time for each image). While illumination intensity cannot reduce spatial noise (since it is not varying with time), the illumination intensity must be increased to the level at which the net tumor signal is above the electronic noise of the detector. However, illumination intensity cannot be arbitrarily increased as limits exist either due to safety requirements or photobleaching of organic fluorophores representing an upper limit to illumination intensity and duration with estimates that each fluorophore can repeat the excitation-emission cycle 10,000-40,000 times before permanently photobleaching ^26^.

### Fluorophore Quantum Efficiency

Photons incident on a labeled cell interact with the bound fluorophore to produce a lower energy, Stokes shifted, fluorescently emitted photon. The efficiency of this process is directly proportional to the optical signal intensity. However, the relative low efficiency of this process requires illumination intensities 3 to 6 orders of magnitude larger than the fluorescence emission. This is driven by the small fluorophore absorption cross section, on the order of 10^−16^cm^2 27^ which defines the area in which a photon can interact with a fluorophore, and the fluorescence quantum yield, which quantifies the probability that a photon interacting with a fluorophore will produce an emitted photon. The quantum yield is typically between 3-10% ^28,29^ for organic fluorophores used in *in vivo* applications.

### Electronic Detection (Responsivity)

Each pixel converts incident photons to electrons, which in turn are converted to a voltage for electronic readout. Each pixel’s total capacity for integrated electrons is the sum of the signal and total background with electronic noise. Pixel responsivity quantifies the efficiency of converting received photons into electrons, which can then be converted to a voltage for electronic readout.

### Pixel Size

The maximal signal in a pixel can be obtained by matching the area imaged to the area subtended by the tumor cells being imaged. This optimizes signal detection as pixel capacity is not consumed by background photons from neighboring normal tissue. As an example, with a goal of imaging 100 cells, this area is approximately 15,000 μm^2^ or roughly a 120 μm x 120 μm for a 10 μm x 10 μm x 10 μm cell. However, maximizing signal in this manner comes with the sharp tradeoff of resolution, which can compromise performance of automated image recognition and machine learning algorithms.

### Quantification of Background

The background is comprised of electrons integrated by the pixel from sources other than the labeled tumor cells. Given all electrons are identical at the pixel level, electrons generated from signal and background are indistinguishable. Therefore, background must be accurately subtracted from the total pixel signal to yield the tumor signal. Here we describe the electronic, optical, and biological sources of background: dark current, optical bleed through, autofluorescence, on-target off tumor binding, as well as non-specific binding.

### Dark Current

Electronic sources of background are primarily due to the dark current ^30–32^. The magnitude of the dark current is dependent on the process used to manufacture the sensor and the sensor operating temperature. While assumed to be constant across pixels, fabrication miss-match between pixels and subtle integrated circuit fabrication process variations result in pixel-to-pixel variation, and this value is best measured per pixel (in darkness), and subtracted from the final readout. The relative contribution of this source of noise can be decreased through longer integration times or averaging multiple images.

### Optical Bleed Through

Poor fluorophore efficiency necessitates illumination intensities orders of magnitude greater than the emitted light. Identifying fluorescently labeled cells thereby requires high performance optical filters that can reject light differing by ~50 nm by 4-6 orders of magnitude. These filters inevitably allow some light through contributing to background, consuming pixel capacity, and increasing shot noise contributions.

### Autofluorescence

Autofluorescence results from a broad-spectrum optical emission of higher wavelength light by molecules in tissue when excited by light. A portion of this emission falls within the emission band of the fluorophore (and therefore the selected optical filter), and as such, is imaged along with the tumor signal. While autofluorescence is reduced using NIR illumination light, it represents a significant source of background as each cell (both tumor and normal tissue) contribute to this background signal.

### On-target, off tumor labeling

Healthy (eg non-cancerous) tissue cells often express a baseline amount of biomarker which binds to the optical label and contributes directly to the background signal. This background is particularly problematic because it appears identical to the tumor signal, as both emit at the same wavelength and cannot be blocked by the filter.

### Non-Specific Binding

Imaging agent adhering to cells that do not express the surface marker or are not eliminated from the patient directly contribute to background. This non-specific binding limits pixel capacity for signal and is a major hindrance for optical imaging of microscopic disease. This is addressed through increased pixel capacity to accommodate the additional background light. The penetration of light into tissue (as with NIR illumination) further adds to this as non-specific binding to cells below the surface also contribute to background. This background can be reduced with a lower wavelength fluorophore, although this sacrifices penetration of superficial, overlying layers of blood that may be found intraoperatively.

## Noise

Various sources of biochemical, physical, optical, and electronic features contribute to variation in the background, obfuscating the appropriate signal for background subtraction. Broadly speaking, electronic noise varies over time, and therefore can be reduced by averaging multiple images. However spatial noise is a function of the tissue itself, and does not change with time (at least within the short interval of intraoperative imaging), and therefore cannot be reduced with averaging. Here we describe biochemical and physical sources of noise as forms of spatial noise.

### Time Varying Electronic Noise

Shot noise represents the fundamental physical limit of detection of counting electrons (generated from incident photons), including all sources of electrons such as optical background and dark current. Consequently, in the presence of a significant background signal, even if accurate background subtraction can be ensured, the noise from a large background signal (but not necessarily the background signal itself) can mask a small signal due to this noise source alone.

### Spatial Noise

The relative ease of subtracting a constant background would obviate the need for SNR considerations. However, background cannot be measured to an arbitrary precision and variation can inhibit our ability to identify the signal of interest. This background variation can occur over a wide frequency scale. Notably, variations in antibody distribution and binding cannot be predicted *a priori*, prohibiting the use of a global threshold for MRD and current efforts at background subtraction are limited to centimeter-scale tumor foci ^9^. For example, the standard deviation for antibody binding per square micron in tissue labeled with an antibody *in vivo* (Figure 2) ranged from 1,259 antibodies/μm^2^ in a HER2-negative tumor (e.g. background variation) to 21,807 antibodies/μm^2^ in HER2-overexpressing cells (e.g. tumor variation and heterogenity). This drives the need for more accurate, patient (and tumor) specific background measurements.

Spatial noise is divided into high frequency (e.g. rapidly varying) spatial noise and low frequency spatial (e.g. slowly varying) noise wherein the high frequency component is composed of cell to cell binding variations or tissue heterogeneity and the low frequency component is composed of gradients in the signal due to diffusion of the antibody. Both high and low frequency variations inhibit the ability to identify an accurate background to subtract and drive the need to quantify the optical pixel size for background subtraction. High frequency spatial noise can be addressed through averaging a larger area - driving the need for a larger pixel size, while low frequency spatial noise precludes too large a pixel size to avoid integrating slowly varying intensity changes over the tissue surface. Thus there is an optimal pixel size to minimize noise, and maximize signal.

#### Noise Quantification

To quantify both the high and low frequency noise we measured the variability of antibody staining across a tissue slice as a function of position on the slide. This tissue slice of an 8 mm HER2+ tumor, shown at the bottom of Figure 4, was resampled at various simulated pixel sizes to demonstrate the impact of noise at different resolutions. At a small pixel size (5 µm, Figure 4, dotted blue trace) high frequency spatial noise fluctuations render local background identification impossible. At large pixel sizes (500 µm, Figure 4, yellow trace), low frequency drift also impairs background identification in a focal area.

**Figure 4.**
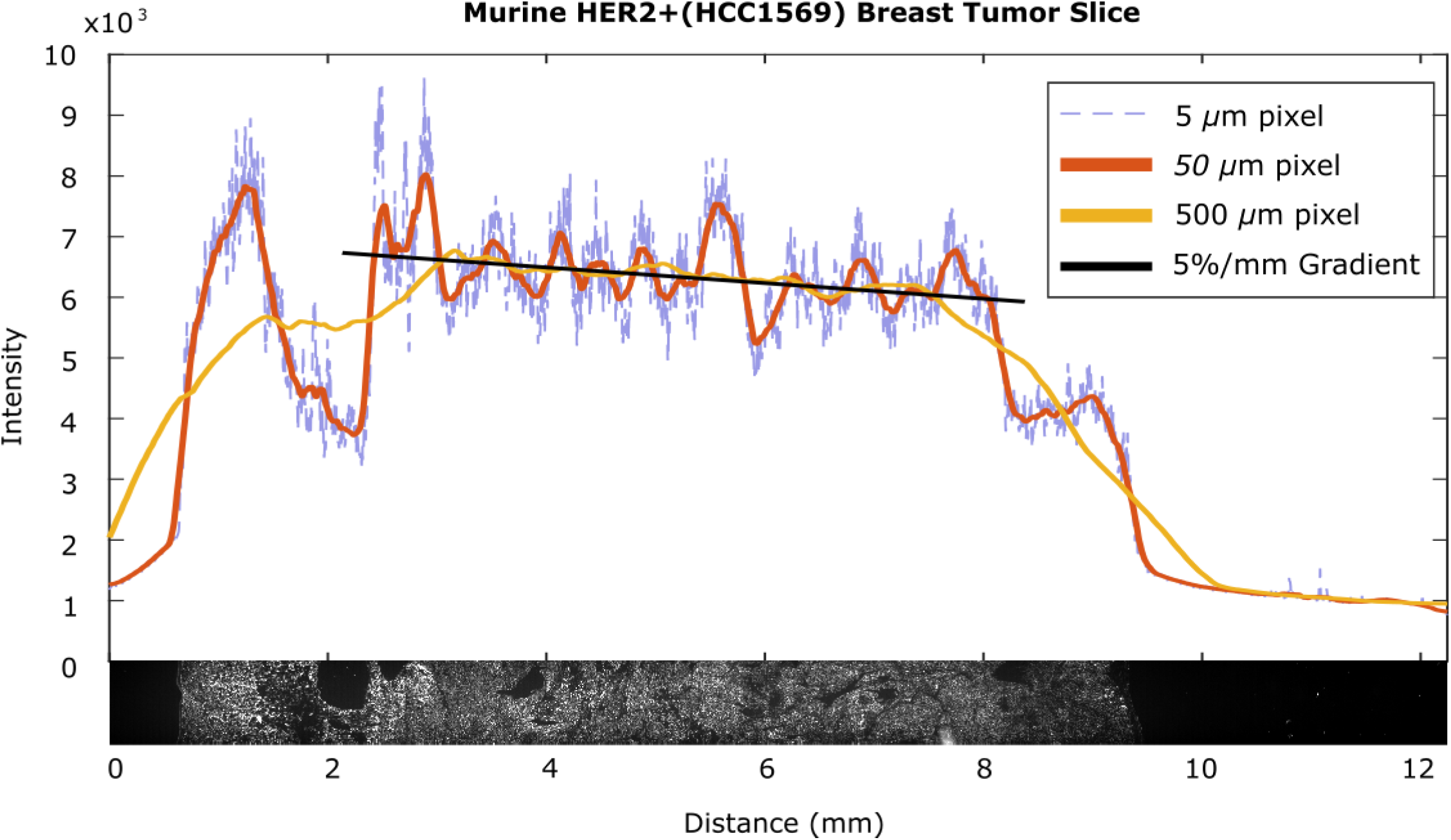
Low frequency spatial variations in a tumor slice. A section of HER2+ (HCC1569) tumor alongside line scans at various spatial resolutions demonstrates the heterogeneities that make determining background difficult. The 5 μm pixel information demonstrates high frequency variation between cells, the 50 μm pixel information demonstrates the background variation due to tissue physiology and the linear portion of the 500 μm pixel information between 3 mm to 7 mm demonstrates a 5%/mm gradient likely due to diffusion

### Electronic Noise

In this example, we assume that adequate averaging reduces electronic noise to below the noise level of the spatial noise, and disregard it.

### Low Frequency Spatial Noise

When sampling background at large distances from the tumor the background changes as a function of the distance from the tumor cluster being imaged, and can be thought of as “low frequency” spatial noise. For example in Figure 4 (yellow trace), this can be as large as 5%/mm when sampling with a 500 μm pixel. This drives the need for high spatial resolution, so that a background measurement can be taken from a pixel close to the pixel imaging microscopic tumor, reducing this noise component.

### High Frequency Spatial Noise

However, high frequency variation puts an upper limit on the spatial resolution: as seen in Figure 4 (blue dashed trace), sampling using 5 μm pixels shows marked pixel-to-pixel variation due to cell to cell variations which can be quantified as the standard deviation in background. Hence, the optimal spatial resolution must sufficiently average the cell to cell variation without merely detecting the drift in intensity due to low frequency noise effects such as antibody diffusion, justifying measurement of these variables for imaging of microscopic disease.

#### SNR Calculation

To calculate the spatial noise as a function of pixel size, we find the variance across the pixels in the image by finding the mean of the square of differences between neighboring pixels as follows. If P_x,y_ is the intensity of pixel at position x,y and N_pixel_ is the number of pixels on the sensor, then noise ΔB is as follows

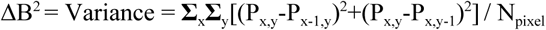

Variance is the more relevant measure of noise to determine SNR. To demonstrate this, we compute the variance using our metric at various spatial resolutions for the image in Figure 1, and plot the results in Figure 5. Below this plot, we show the image as it appears at given spatial resolutions (a-e). Variance across the image changes as a function of pixel size with the expected behavior of decreasing variance as high frequency noise components are reduced with increasing spatial averaging, until low frequency noise dominates and the variance begins to increase again. Thus an intermediate pixel size (“c”) provides the optimal SNR for this system.

**Figure 5.**
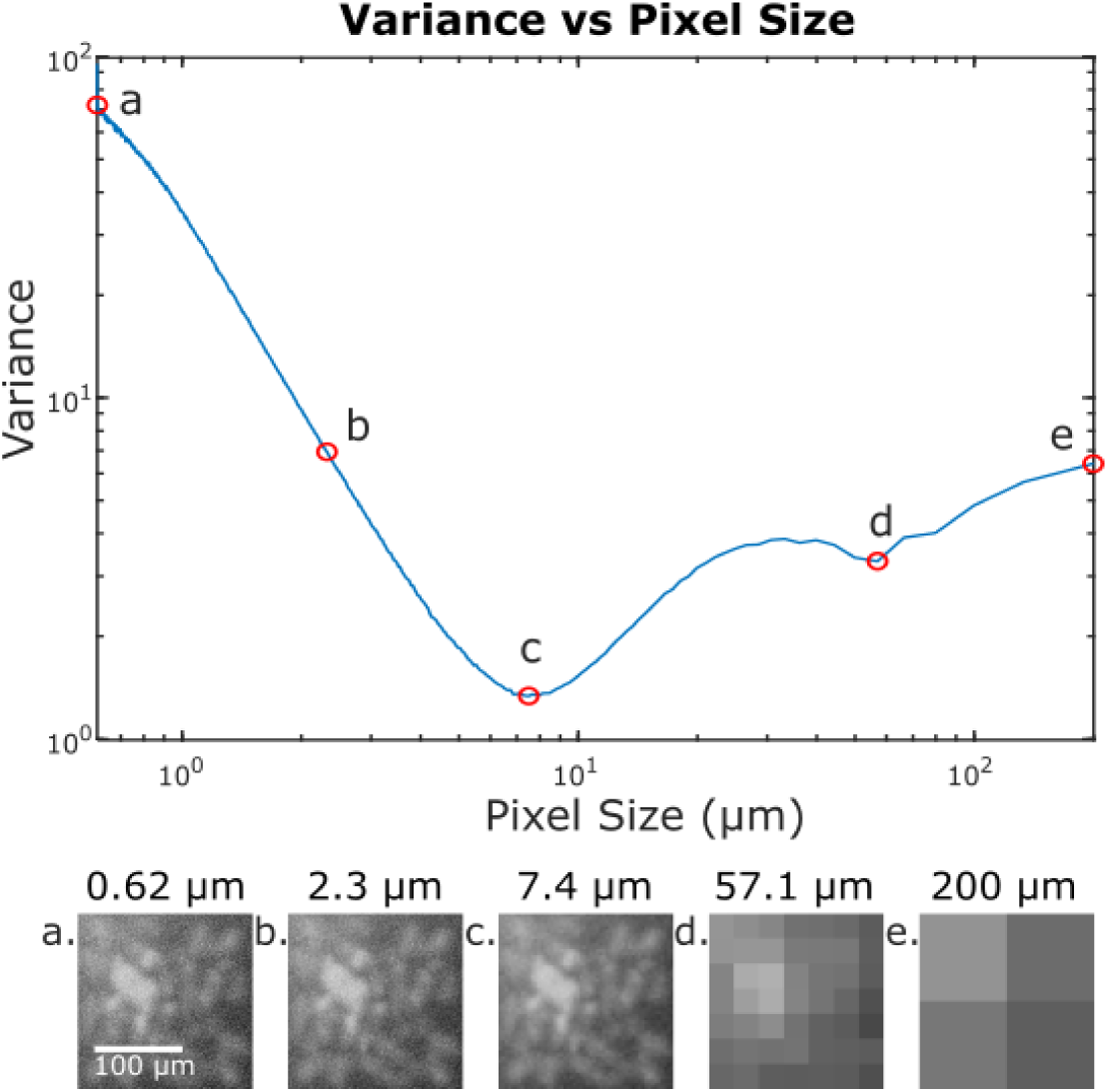
Variance in the image from Figure 1 at varying spatial resolutions. **(a)** represents the original image (**b)** illustrates a reduction in high frequency noise without significant loss in resolution. The minimum variance across the image occurs at point (**c)** at a pixel size that averages the high frequency noise component without being dominated by low frequency noise. Point (**d)** has increased variance due to averaging of low frequency noise components and in point **(e)**, this low frequency noise is the only feature visible.

#### Monte Carlo Simulation of Maximal SNR and Optimal Pixel Size

To ensure that our results are generalizable and not simply driven by our chosen sample, we ran a monte carlo simulation of 50 computer generated cell images. Each image consists of randomly generated cell clusters with an average of 100 cells, with signal, background, and spatial noise derived from measured image data as described.

In Figure 6, we plot the signal to noise ratio over spatial resolutions corresponding to pixel sizes ranging from 0.61 μm to 200 μm for 50 random cell clusters. One of these random clusters is shown at select resolutions to illustrate how optimized SNR can enhance micro-tumor identification. In this instance, the optimal pixel size is in the range of 10 μm - 35 μm with a maximum SNR of 25.

**Figure 6.**
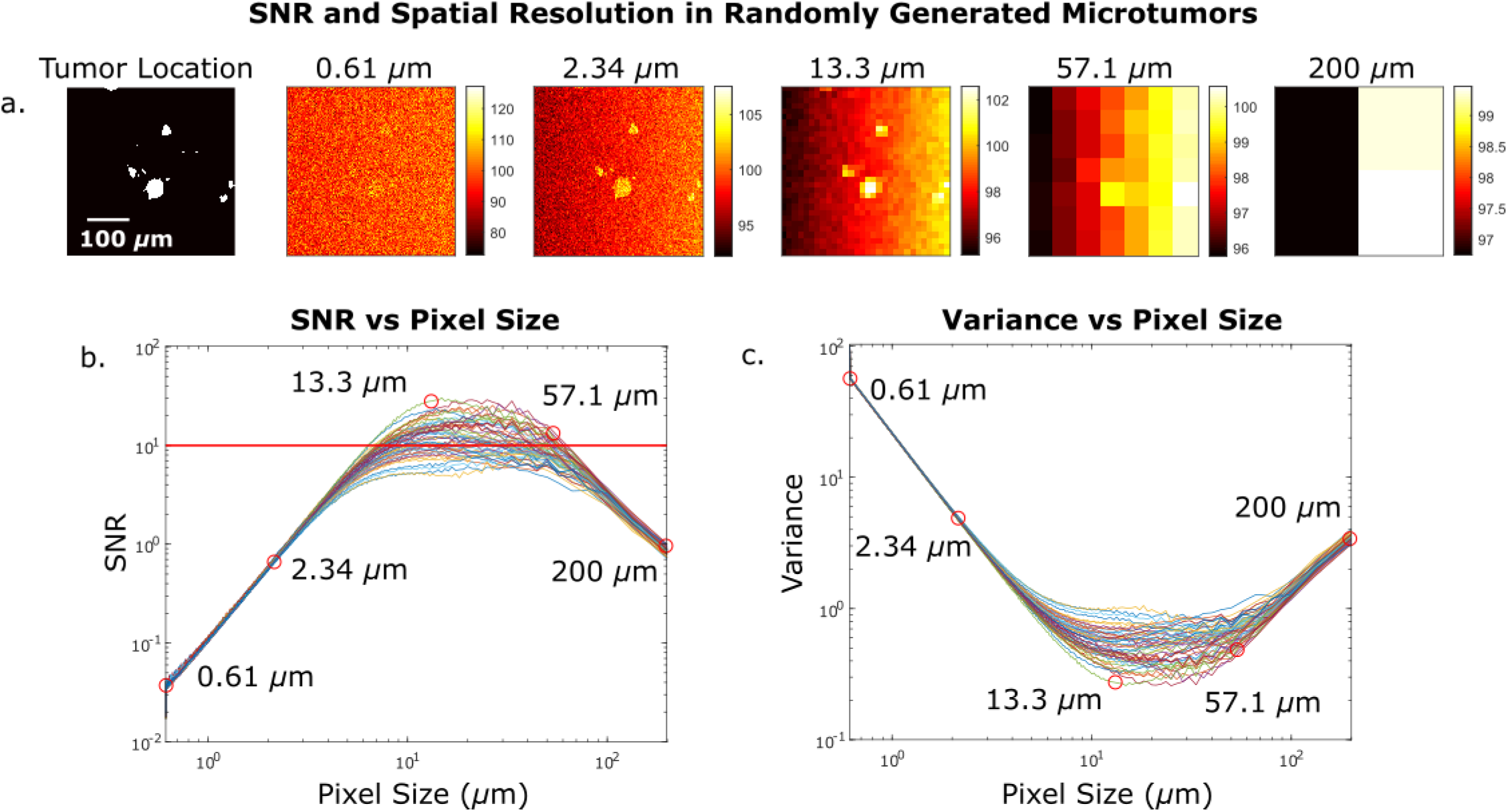
Monte carlo simulation of procedurally generated tumor images. **(a)** Example of randomly generated tumor followed by images at various spatial resolutions with simulated noise. The average background and noise are modelled after empirically determined distributions. **(b)** Plot of SNR at various pixel sizes over 50 randomly generated tumors, red line demarcates SNR = 10. **(c)** Variance across an image at various pixel sizes. Variance decreases as high frequency noise is averaged away, then increases as low frequency noise dominates.

#### Extension to Single Cell Imaging

Assuming a sparse distribution of tumor cells, a single cell with sufficient SNR can be detected by a pixels much larger than the size of the cell. We simulated a 10μm cell with tumor to background ratios from 1-30 to cover the real world range we found in *in vivo* staining data shown in Figure 3. In Figure 7, we present an instance of this simulation with images of a single cancer cell at the center of a background of healthy tissue. This cell has a TBR of 10.4 while the healthy cells have random intensity with the same mean intensity and distribution as empirically determined. Even pixels an order of magnitude larger than the cell can uniquely locate a single cancerous cell, however there is an upper bound to pixel sizes used to locate single cells. With a SNR below 10, there is an increasing chance that other pixels yield a greater intensity than the pixel over the tumor cell, rendering unique identification impossible. Establishing that the maximum size of the pixel that can reliably detect a single cell occurs when the SNR is greater than 10, we plot the SNR at various spatial resolutions for the example TBR of 10.4. In addition, we plot the largest pixel size that achieves an SNR of 10 for the full range of TBRs. Figure 7 illustrates that single cell imaging is achievable even in an optical imaging system with resolution lower than that of a single cell.

**Figure 7.**
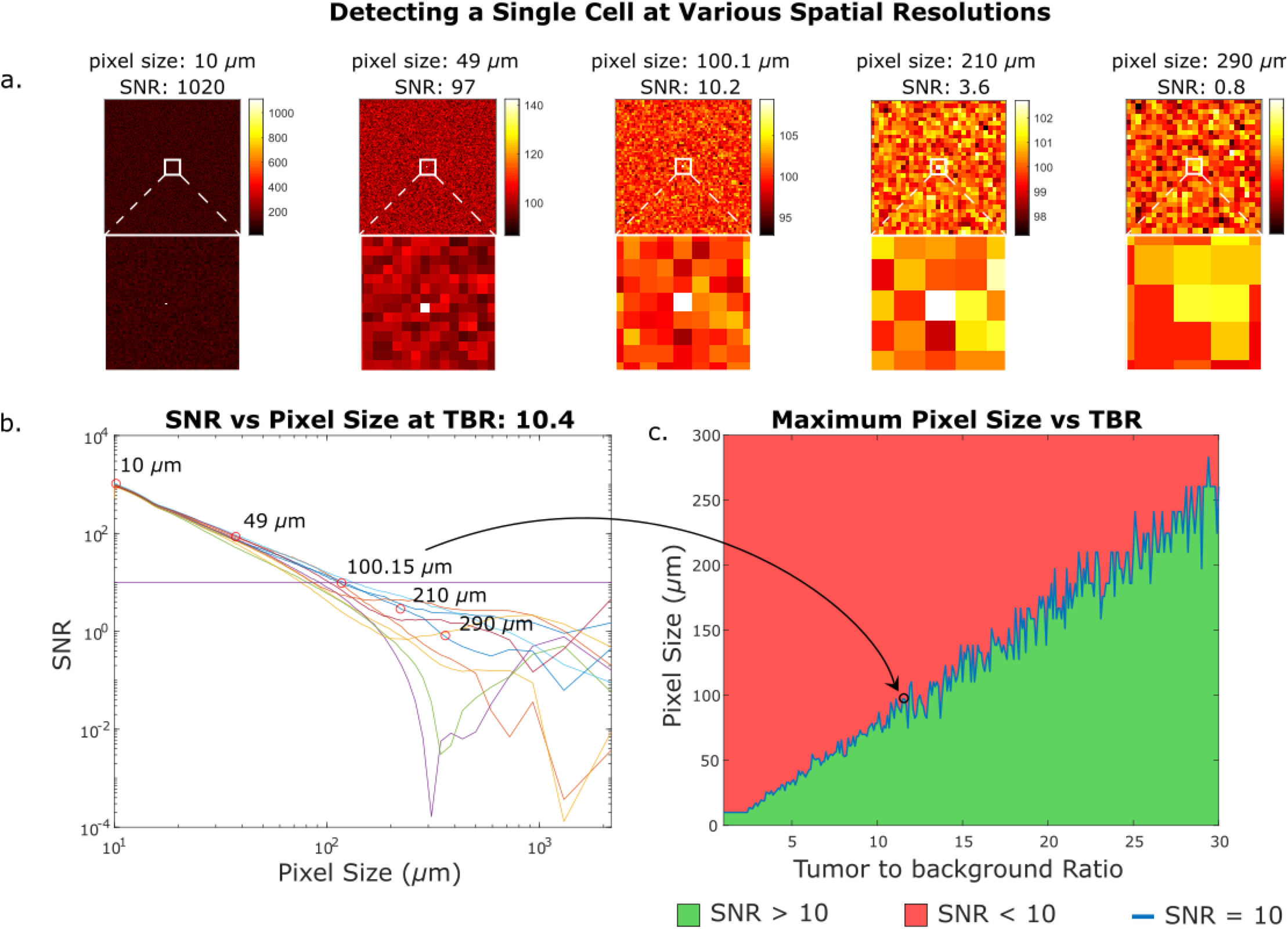
Detecting a single cell at various spatial resolutions. In **(a)** we see the results of imaging a 10 micron diameter cell in healthy tissue with some noise and a tumor to background ratio of 10.4. We can still determine the location of the cell at the center of the imager as long as the SNR is greater than 10. Detecting a single cell does not require sub-cellular resolution contingent on the notion that the single cells are sparsely distributed and have a large tumor to background ratio. In **(b)** we see the SNR over a range of pixel sizes for 10 randomly generated samples given a TBR of 10.4. In **(c)** we plot the largest pixel that can detect a single cell at a given TBR with SNR greater than 10. The plot is not smooth because it is based on randomly generated samples with arbitrary noise.

## Discussion

In this study we have outlined and demonstrated a method to characterize the ability of optical imagers and targeted molecular imaging agents (TMIAs) to identify microscopic tumor foci, including single cell residual disease. These small areas of tumor often exhibit intensity on the order of background, necessitating a metric beyond signal to background ratio. Furthermore, the advent of machine learning and automated image recognition algorithms lend themselves to a more quantitative evaluation of imaging system performance. Through characterization of the signal intensity per cell, background intensity per cell, and the variation in background, the ultimate level of sensitivity for a given foci of tumor cells can be calculated. Furthermore, we demonstrate a methodology for simulating imaging of microscopic disease using computer generated images, allowing evaluation of the sensitivity of an imaging system and companion TMIAs.

We can use this analysis to optimize the design parameters such as pixel size for future intraoperative imagers. The current paradigm is to pursue small pixels for high resolution images. However, while higher resolution images can be binned to create a larger pixel size in post-processing software, higher pixel density has intrinsic costs: smaller pixels have relatively more temporal nose (as they integrate less signal), requiring longer averages; the fill factor is reduced (for example CMOS-based imagers often include in-pixel electronics); and longer readout times are necessary to obtain the data from more pixels. Therefore, it is advantageous to have the optimal sized pixel within the system itself.

Pixels cannot be arbitrarily large either, both to retain resolution for accurate location data, and to prevent capturing low frequency spatial noise across the image.To image microtumor with ~100 μm diameter and background noise consistent with trastuzumab or J591 labelling, pixels sizes between 10 μm −35 μm yielded optimal results (Figure 6). As pixel size increases up to 10 μm, the variance across the image decreases, while pixels over the tumor capture a greater portion of the signal, causing an increase in SNR. As pixel size continues to increase past the optimal range, the image comes to be dominated by any underlying diffusion or tissue patterns that mask the tumor location.

In Figure 6, we see tumor location is not readily visible due to high frequency noise with a small pixel size of 0.61 μm. Spatial averaging by increasing pixel size to 2.34 μm results in improved SNR and tumor areas are better defined. However, there is still large variance across the image that may limit automatic detection. Peak SNR with sampling at 13.3 μm pixel size yields clear identification of microtumor areas. Tumor areas are still identifiable even at low resolution with a pixel size of 57.1 μm. High frequency spatial noise is reduced until low frequency spatial noise dominates at larger pixels such as 200 μm where only the gradient is visible. Of particular note, a low pixel size may result in easily identifiable tumor areas, such is the case at a pixel size of 2.34 μm. However thresholding for automatic tumor detection would not work well as there are many background pixels with high intensities outside tumor areas. Increased spatial averaging, either in post processing, or at a hardware level with larger pixels rectifies this high noise.

In the extreme condition of detecting a single tumor cell amongst a background of non-cancerous tissue, detectors with pixels ranging from 10 μm - 250 μm can be used corresponding to TBRs ranging from 1-30 (Figure 7c). If a detector has pixels that are too large, then background areas distant from the tumor cell may have average intensities that appear to be tumor, as seen in Figure 7a with a 210 μm pixel size. This results in a degeneration of SNR where the area with a tumor cell cannot be uniquely identified.

While the metrics incorporated here address the image quality and ability to identify microscopic disease with optical imagers, they do not address other key metrics of intraoperative imagers including imager size, mobility, and ability to fit within hard to access areas allowing visualization of all sides of a tumor cavity and within lymph node basins. For example, fiber optics have a fundamental tradeoff between fiber diameter (which directly relates to the area visualized with each image) and flexibility, with a 1 cm bending radius achievable only with optical fibers of roughly 100 μm diameter. Similarly, imaging speed is important, as the entire surface area must be imaged rapidly to enable seamless integration into surgery and prevent the image from being degraded with hand motion.

Imaging small numbers of tumor cells with very low fluorescence levels has a large impact for guiding cancer surgery and requires the assistance of image processing algorithms (Carpenter et al. 2006). These techniques can be used synergistically with intraoperative imagers designed to image broad areas ^33^ and guide gross resection. Following initial removal, tools adept at quantification and characterization of MRD can assist in the decision to further resect with the goal of achieving negative margins.

## Conclusion

Detecting and removing microscopic disease in margins during tumor resection has significant impact on patient care and outcomes. A growing array of sensors, imagers, and optical labels address this problem of intraoperative imaging and are largely characterized by the tumor to background ratio they can detect as a proxy for human detection. Here we show that the spatial signal to noise ratio (SNR) is a fundamental limit of electronic image detection and describe techniques to quantify signal and noise in image systems as well as optimizations to improve SNR for the most accurate detection. We demonstrate our results using a monte carlo simulation of SNR in procedurally generated tumor images based on parameters of signal, background, and noise that we quantified from imaging HER2-overexpressing and HER2-negative cell lines with fluorescently labelled trastuzumab and PSMA-positive and PSMA-negative cell lines with fluorescently labeled J591 antibody. We extend this SNR analysis to optical imaging systems for single cell detection.

## Acknowledgements

MA was supported by NIH R21EB027238 and DOD PC141609. This project is supported by DOD PC141609. We would like to thank Hui Zhang for performing the *in vitro* and *in vivo* antibody staining.

